# Learning of pathogenic bacteria in adult *C. elegans* bidirectionally regulates pathogen response in the progeny

**DOI:** 10.1101/500264

**Authors:** Ana Pereira, Xicotencatl Gracida, Konstantinos Kagias, Yun Zhang

## Abstract

Parental experience can generate adaptive changes in the behavioral and physiological traits of the offspring^1–3^. However, the biological properties of this intergenerational regulation and the underlying molecular and cellular mechanism are not well understood. Here, we show that the experience of learning to avoid pathogenic bacteria in *C. elegans* alters the behavioral response to the pathogen in the progeny through the endogenous RNA interference (RNAi) pathway. We previously show that the adult *C. elegans* learns to avoid the smell of pathogenic bacteria, such as the *Pseudomonas aeruginosa* strain PA14, after feeding on the pathogen for a few hours^4,5^. Here, we report that this learning experience can bidirectionally regulate the olfactory response to PA14 in the progeny that are never directly exposed to the pathogen. The olfactory preference for PA14 in these progeny is linearly correlated with the learned avoidance of PA14 in their mothers. If the mothers show strong learning of PA14, their progeny avoid PA14; intriguingly, if the mothers show weak learning of PA14, the progeny prefer PA14, suggesting that the PA14-trained mothers transmit both the negative and positive information of PA14 to their progeny. The intergenerational behavioral effect results from an altered behavioral decision regulated by an olfactory sensorimotor neural circuit. Learning to avoid the pathogen also influences the development of the progeny, which is regulated independently from the behavioral change. Animals mutated for the RRF-3/RNA-directed RNA polymerase, a master regulator for the synthesis of the small interfering RNAs that are maternally inherited or in the soma^6,7^, display the normal naive and learned response to PA14 but are defective in regulating the olfactory response to PA14 in their progeny. Our results characterize an intergenerational effect that allows the progeny to rapidly adapt to an environmental condition that is critical for survival.

---

In many organisms, the parental experience generates adaptive changes in the behavior and the physiology of the progeny^1–3^. However, whether the parental interaction with the environment regulates the same behavioral modalities in the parents and the progeny, indicating the transmission of specific sensory information, is not clear. In addition, the signaling pathways underlying the transmission of the parental information to the progeny are poorly understood. Olfaction plays critical roles in animal behaviors that are essential for survival, such as searching for food and avoiding dangers. While genetically encoded^8,9^, the olfactory system is highly adaptable. Experience can profoundly shape the meaning of an odorant to an animal^10,11^. *Caenorhabditis elegans* feeds on bacteria and is often attracted by the smell of the bacteria in its habitat^4,12,13^. Meanwhile, some bacteria are pathogenic. Ingesting the pathogenic bacteria makes the worm ill through intestinal infection^14^. Previous studies show that after briefly feeding on certain pathogenic bacteria, such as the *Pseudomonas aeruginosa* strain PA14, adult *C. elegans* reduces the preference for the smell of the pathogen, even if it is initially attractive^4,5^. This form of learning is contingent on the pathogenesis of the training bacteria^4,5^ and resembles the Garcia effect, a robust form of conditioned aversion that allows animals to learn to avoid the smell or taste of a food that makes them ill^15^.

Although the intestinal infection caused by ingesting PA14 results in a slow death of the worm over a course of few days^14^, the bacteria in the *Pseudomonas* genus are abundant in the natural habit of *C. elegans* and serve as common food sources for the worm^13^. Therefore, the pathogenic *Pseudomonas* PA14 contains the information of food and danger, both of which are critical for the survival of the worm. We asked whether learning to avoid PA14 regulated the olfactory response to the smell of PA14 in the offspring. We cultivated the worms under the standard conditions on the benign bacteria strain *Escherichia Coli* OP50 until the adult stage^16^. We then trained the adult hermaphrodite worms (F0s) by feeding them on PA14 for 8 hours (Figure 1A). We quantified the preference between the smell of OP50 and the smell of PA14 in the trained and the naive F0 worms using an automated assay^5^. In this assay, a positive choice index indicates a preference for PA14 and a negative choice index indicates a preference for OP50 (Methods)^5^. After randomly sampling the preference in F0s, we harvested the embryos (F1s) from the trained and the naive F0s with a bleach solution that dissolved the adult bodies and the associated bacteria. We cultivated the F1 worms on the benign bacteria *E. Coli* OP50 under the standard conditions. Thus, the F1 progeny of the PA14-trained F0s and the F1 progeny of the naive F0s were cultivated under the same condition and neither was directly exposed to PA14. We quantified the preference of F1s between the smell of OP50 and the smell of PA14 after they reached the adult stage. We measured the olfactory preference of F1 worms using a two-choice assay, in which a worm chooses between a drop of supernatant of an OP50 bacteria culture and a drop of supernatant of an PA14 bacteria culture by navigating towards the drop (Figure 1A). We recorded the behavior of the F1 worms during the two-choice assay and analyzed their chemotactic movement and their choices. A positive choice index in the two-choice assay indicates a preference for PA14 and a negative choice index indicates a preference for OP50 (Figure 1A and Methods). While the automated assay allows us to rapidly measure the olfactory preference in F0s before handling the F1 embryos, the two-choice assay allows us to examine the olfactory behavior in F1 worms in detail (Methods).

**Figure 1.**
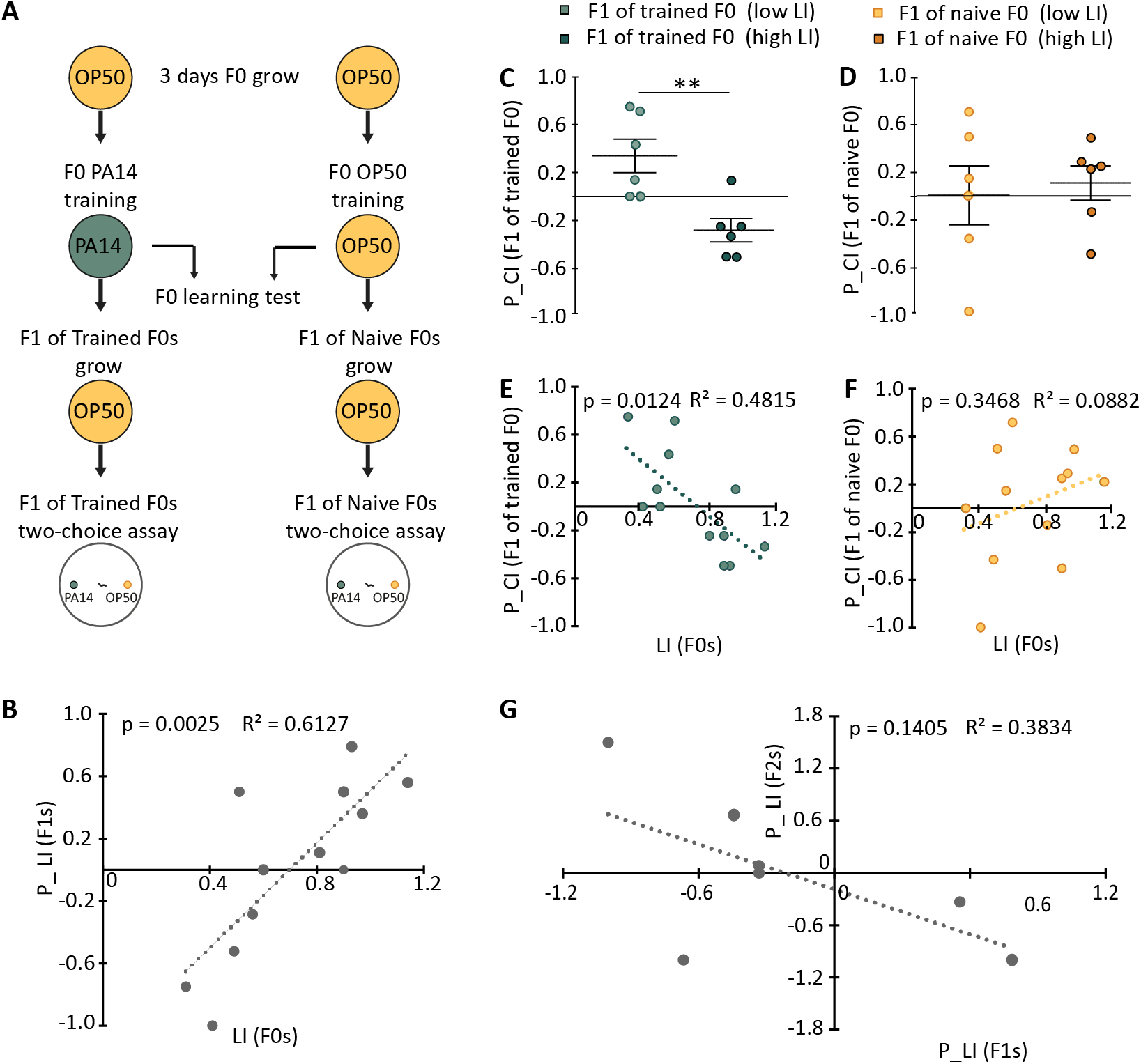
Training with PA14 bidirectionally regulates the olfactory response to PA14 in the progeny. **(A)** Schematics for the parental-experience dependent olfactory learning. **(B)** Parental-experience dependent olfactory learning indexes (P_LI) in F1s are linearly correlated with the aversive olfactory learning indexes (LI) of the F0 mothers. Choice index of F0 (CI) = (bends evoked by OP50 - bends evoked by PA14)/(bends evoked by OP50 + bends evoked by PA14); LI = CI of naive F0s – CI of trained F0s; Choice index of F1 (P_CI) = (number of worms that choose PA14 - number of worms that choose OP50)/total number of worms. P_LI = P_CI of F1s of naive F0s – P_CI of F1s of trained F0s; n = 12 independent experiments that contained 12 independent learning assays for F0s (274 animals) and 12 independent two-choice assays where 186 F1 worms were individually tested, p value denotes the significance of the linear correlation coefficient. **(C, D)** The F1s of the trained F0 mothers in experiments in which F0s exhibited a high level of aversive learning of PA14 avoided PA14, indicated by a negative choice index (P_CI), and the F1s of the trained F0 mothers in experiments in which F0s exhibited a low level of aversive learning of PA14 preferred PA14, indicated by a positive choice index (P_CI) **(C)**; in contrast, the F1s of the naive F0s tested in the same sets of experiments as shown in **C** where F0s exhibited a high level or a low level of learning had similar choice indexes (P_CI) **(D)**. n = 6 experiments for high F0 learning and 6 experiments for low F0 learning, where 95 (high F0 learning) and 91 (low F0 learning) F1 worms individually tested in the two-choice assay for each group; Student’s t-test (data are normally distributed), ** p = 0.004 for **C** and p = 0.72 for **D**; Mean ± SEM. **(E, F)** The choice indexes of the F1s (P_CI) of the trained F0s are inversely correlated with the learning indexes (LI) of their trained F0 mothers **(E)**; in contrast, the choice indexes of the F1s (P_CI) of the naive F0s have no correlation with the learning indexes (LI) of their naive F0 mothers **(F)**. A positive or a negative P_CI respectively indicates a preference for PA14 or OP50, n = 12 learning assays for F0s (n = 274 animals) and n = 12 two-choice assays where 186 F1 worms were individually tested, P values denote the significance of the linear correlation coefficients. **(G)** Parental-experience dependent olfactory learning indexes (P-LI) in F2s is not significantly correlated with the P-LI of the F1s. P-LI = choice index of F1s (or F2) of naive F0s – choice index of F1s (or F2) of trained F0s; choice index of F1 (or F2) = (number of worms that choose PA14 - number of worms that choose OP50)/total number of animals. n = 6 independent experiments that included 91 F1 worms and 88 F2 worms individually tested in the two-choice assays, P values denote the significance of the linear correlation coefficient.

We found that consistent with our previous findings^4,5^, training the F0 adult worms with PA14 for 8 hours strongly reduced the preference for PA14 (Extended Figure 1). This experience-dependent change is indicated by a <u>l</u>earning <u>i</u>ndex (LI), which is the difference between the choice index (CI) of the naive F0s and the choice index of the trained F0s in the automated assay (Figure 1B and Methods). A positive LI indicates learned avoidance of PA14 (Methods). Next, we examined whether the training experience of the F0s generated any difference in the olfactory preference of their F1 progenies. Thus, we measured the difference between the choice index of the F1 progeny of the naive F0 worms and the choice index of the F1 progeny of the trained F0 worms, and defined this difference as the <u>p</u>arental-experience dependent <u>l</u>earning <u>i</u>ndex (P_LI) (Figure 1B and Methods). A positive P_LI indicates increased avoidance of PA14 in F1 induced by the parental experience with PA14 (Methods). We found that P_LI in F1s were significantly correlated with the LI in their F0 mothers (Figure 1B). On one end of this spectrum, the more the trained F0s learned to reduce their preference for the PA14 smell, the more their F1 progeny avoid the PA14 smell in comparison with the F1s of the naive F0s that were tested in parallel (Figure 1B), indicating that a strong aversive learning of PA14 in the F0s generates the avoidance of the PA14 smell in F1. Surprisingly, on the other end of the spectrum, we found that for F0s that only weakly learned to reduce their preference for the PA14 smell, the less the trained F0s learned to suppress their preference for PA14, the more their F1s preferred the PA14 smell in comparison with the F1s of the naive F0s tested in parallel (Figure 1B), suggesting that the F0s that weakly learned transmit the beneficial information of PA14. These results show that the maternal learning experience with PA14 bidirectionally regulates the olfactory response to PA14 in the progeny.

Next, we further examined the bidirectional effect of the F0’s learning experience on the olfactory choice of F1s. We separated all the experiments into two groups, the experiments in which F0s displayed a high level of learning of PA14 and the experiments in which F0s displayed a low level of learning of PA14 (Methods). We found that the F1 progeny of the F0s with strong learning avoided PA14 in the two-choice assay, indicated by their negative choice indexes (P_CI, Figure 1C). In contrast, the F1 progeny of the F0s with weak learning preferred PA14 in the two-choice assay, indicated by their positive choice indexes (P_CI, Figure 1C). The extent of the F0 mothers’ learning significantly regulated the preference in their F1 progeny. This regulatory effect is also evident in a significant correlation between the learning indexes of F0s (LIs) and the choice indexes of the F1s (P_CIs) of the trained F0s (Figure 1E). In comparison, the choice indexes of the F1s (P_CIs) of the naive F0 mothers in the experiments where training generated strong aversive learning were similar to the P_CIs of the F1s of the naive F0 mothers in the experiments where training generated weak aversive learning (Figure 1D). Consistently, there is no correlation between the choice indexes of the F1s (P_CI) of the naive F0s with the learning indexes of the F0 generation (LIs) (Figure 1F). Thus, it is conceivable that the parental experience with PA14 signals two types of information, aversive information such as pathogenesis and appetitive information such as food, and the weight of these information transmitted to the progeny depends on the degree of the parental learning.

To examine whether the difference in the olfactory response to PA14 in the F1 generation passed onto the F2s, we isolated the F2 worms from the F1 mothers and cultivated F2s under the standard conditions on *E. coli* OP50. Similarly, we quantified the parental-experience dependent learning index (P_LI) in F2s, which was defined as the difference between the choice index of the F2s of the naive F0 worms and the choice index of the F2s of the trained F0 worms. We found that the P_LIs of the F2s did not depend on the P_LIs of their F1 mothers (Figure 1G). These results together indicate that the parental experience with PA14 only regulates the olfactory response of their first-generation progeny, leaving the flexibility for F2s to respond to further changes in the environmental conditions.

To further characterize the parental-experience dependent olfactory learning in F1s, we asked what behavioral changes gave rise to their altered olfactory choice. We analyzed the movement of the F1s when they navigated toward OP50 or PA14 during the two-choice assay (Figure 2A). We measured the navigation index, which is defined as the ratio of the radial speed towards the target and the actual traveling speed to quantify the efficiency of the chemotactic movement towards an odorant source^17^(Figure 2B). We divided the assay plate in sections along the axis of the two drops of the bacteria cultures that were 2cm apart and quantified the navigation index for each section in each worm. We analyzed the median value of the navigation indexes for all the worms in each section as a function of the distance away from the target. We found that the navigation index of the F1s of the trained F0s increased when the animals approached the target, indicated by a significant correlation between the navigation index and the distance from the target regardless of whether the worms chose OP50 or PA14 (Figure 2B); in contrast, the navigation index of the F1 worms of the naive F0s did not correlate with the distance to the target (Figure 2B). We also measured the locomotory speed, the distance that the worms travelled and the time spent on travelling before reaching the target, the rate of reorienting movements, the reversals and the large body bends, and did not find any difference between the F1 worms of the trained F0s and the F1 worms of the naive ones (Figure 2C, 2D and Extended Figure 2A-2C). These results identify different strategies for chemotactic movements used by the F1 progeny of the trained mothers versus the F1 progeny of the naive mothers. Previous studies show that the sensorimotor integration of a cholinergic neural circuit regulates the efficiency of chemotactic movements towards odorants, indicated by the navigation index^17^. These results together identify the neuronal substrates for the parental-experience dependent olfactory plasticity in the F1s.

**Figure 2.**
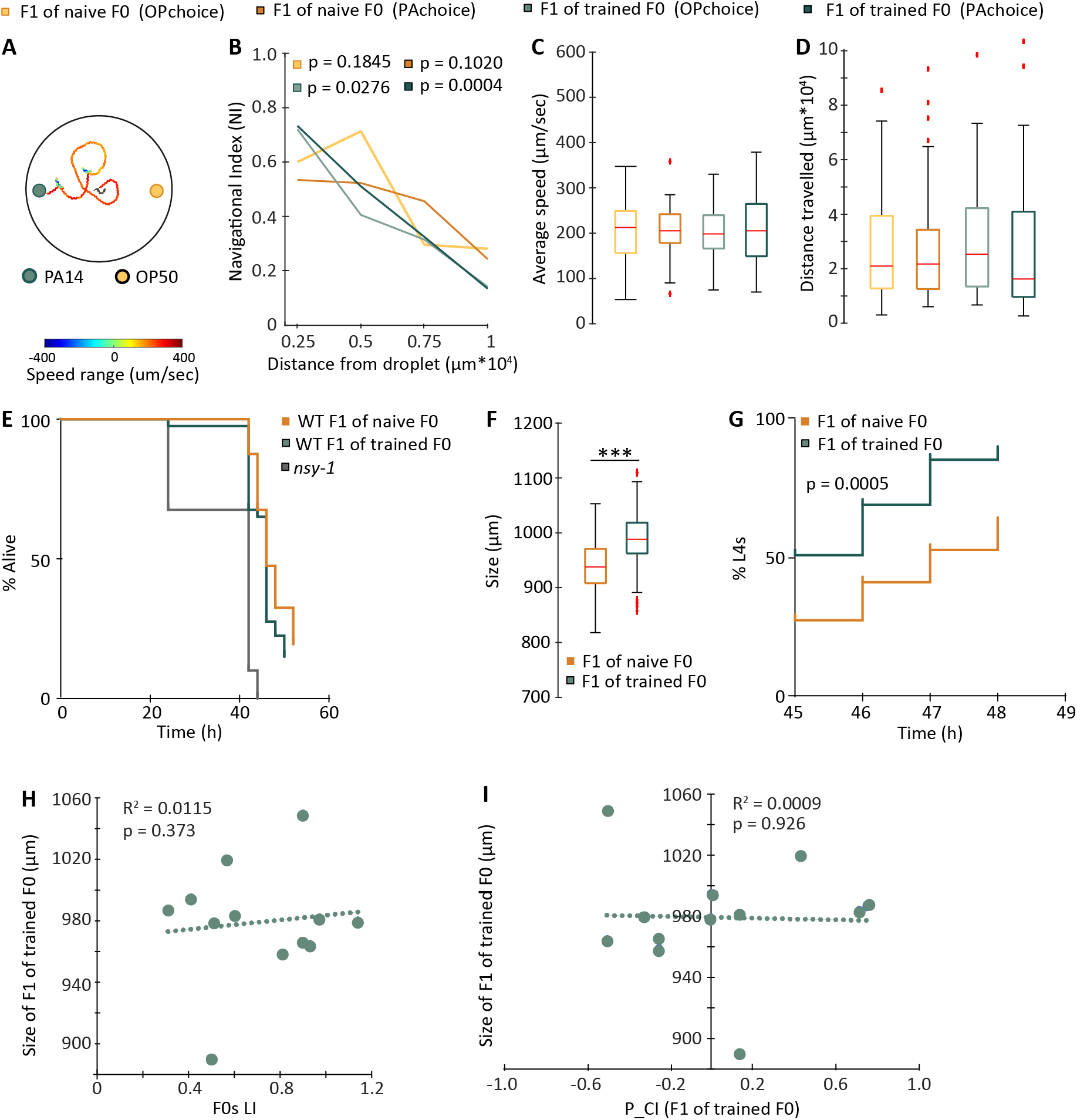
The parental experience with PA14 regulates the olfactory behavior and the development of the F1 progeny. **(A)** Schematics showing the trajectory of a worm in the two-choice assay. **(B)** The navigation indexes in the F1s of the trained F0s inversely and linearly correlate with the distance between the navigating F1s and the target drop of bacteria culture, regardless of the chosen bacteria; in contrast, the navigation indexes in the F1s of the naive F0 mothers do not have the similar correlations. The median value of the navigation indexes for each section was used to analyze linear regression; n = 40 F1s of naive F0 (OPchoice), n = 47 F1s of naive F0 (PAchoice), n = 42 F1s of trained F0 (OPchoice), n = 47 F1s of trained F0 (PAchoice). Data was not analyzed for worms that didn’t made a choice at the end of the trial. Data is missing for 7 worms due to problems during recordings. **(C, D)** The F1s of the trained F0s and the F1s of the naive F0s do not differ in the locomotory speeds during the chemotactic movement towards bacteria cultures **(C)** and traveled similar distances before reaching the target **(D)**. The box plots show the median, the first and the third quantile, 1.5 IQR and the outliers; the number of worms for each group is the same as in **B**, Wilcoxon rank-sum test with correction for multiple comparison (the data is not normally distributed). **(E)** In the slow killing assay, the wild-type (WT) F1s of the trained F0s and the wild-type F1s of the naive F0s did not differ in their survival rates (p = 0.104), while the *nsy-1* mutant animals were significantly weaker in resisting PA14 infection (p < 0.0001). Two independent experiments generated the same conclusion and one experiment containing two replicates (n = 20 animal each plate) was presented, Mantel-Cox log rank test. **(F, G)** The F1s of the trained F0s are larger in the body size than the F1s of the naive F0s [**F**, the box plots show the median, the first and the third quantile, 1.5 IQR and outliers; n = 87 F1s of naive F0 and n = 89 F1s of trained F0, Wilcoxon rank-sum test with correction for multiple comparison (the data is not normally distributed)] and contain a higher percentage of the L4-stage larvae after developing from the embryos extracted from F0s for the same amount of time (**G**, n = 5 experiments with 10 animals for each group in each experiment, Mantel-Cox log rank test). **(H, I)** The body size of the F1s of the trained F0s do not correlate with the learning index (LI) of their F0 mothers **(H)** or their own choice indexes (P_CI) **(I)**. n = 12 experiments in which the learning index of F0s and the choice index of F1s are measured and n = 89 F1 worms were measured for the body size.

In addition, we asked whether the parental experience with PA14 regulated other physiological traits of the F1s. We first examined the immune resistance to PA14 in F1s using a slow-killing assay that measures the survival rate over time in worms cultivated on a lawn of PA14^14^. We used the *nsy-1(k397)* animals as a control, because *nsy-1(k397)* is mutated for the worm homolog of MAPKKK and displays significantly compromised resistance to the infection of PA14^18^. We found that while the *nsy-1(k397)* animals showed a much reduced survival rate, the F1s of the PA14-trained F0s and the F1s of the naive F0s showed similar survival rates (Figure 2E), indicating that training the adult worms with PA14 do not alter the resistance to PA14 in their progeny. However, we noticed that the F1s of the trained mothers were larger in the body size than the F1s of the naive F0 mothers (Figure 2F). To characterize this difference, we examined the F1 embryos inside the F0s. While the trained F0 mothers and the naive F0 mothers contained the same amount of embryos in their bodies on average (trained F0s: 12 ± 0.8 eggs; naive F0s, 11 ± 0.7 eggs; n = 7 adults each, mean ± sem, two-tailed Student’s t-test, p = 0.3), the F1 embryos inside the trained F0s were more advanced in the developmental stage than the F1 embryos inside the naive F0s (Extended Figure 2D). In addition, the F1s of the trained F0 mothers entered the L4-larval stage, a developmental stage with distinct anatomical features^19^, sooner than the F1s of the naive F0 mothers (Figure 2G). Thus, the parental experience with PA14 influences the development of the offspring. Notably, there is no correlation between the body size of the F1 progeny of the trained mothers with either the learning index of the mothers (LI) or the olfactory choice of the F1s (P_CI) (Figure 2H and 2I), indicating that the parental-experience dependent changes in the olfactory choice of the F1s and in the development of the F1s are independently regulated.

Next, we sought the signaling mechanisms whereby the information of the parental experience is transmitted to the progeny. In *C. elegans*, the endogenous RNA interference pathway can respond to environmental conditions to generate small interfering RNAs (endo-siRNAs) that transmit the parental experience cross generations and to regulate the physiology of the offspring^20^. We tested the mutant animals that contained a deletion in *rrf-3*, which encodes a RNA-directed RNA Polymerase(RdRP)^21^ that is needed for the biogenesis of endo-siRNAs that are maternally inherited or found in the soma^6^,^7^. We found that the *rrf-3(pk1426)* mutant animals display normal naive and learned olfactory preference for PA14 in the F0 generation (Figure 3A), but are significantly defective in the parental-experience dependent learning in F1s (Figure 3B). While the parental-experience dependent learning index (P_LI) in F1s significantly correlated with the learning index (LI) in F0s in wild-type animals, this correlation was completely abolished in the *rrf-3(pk1426)* mutants (Figure 3B). Thus, the pathway of the endo-siRNAs regulates the transmission of the parental experience to the progeny.

**Figure 3.**
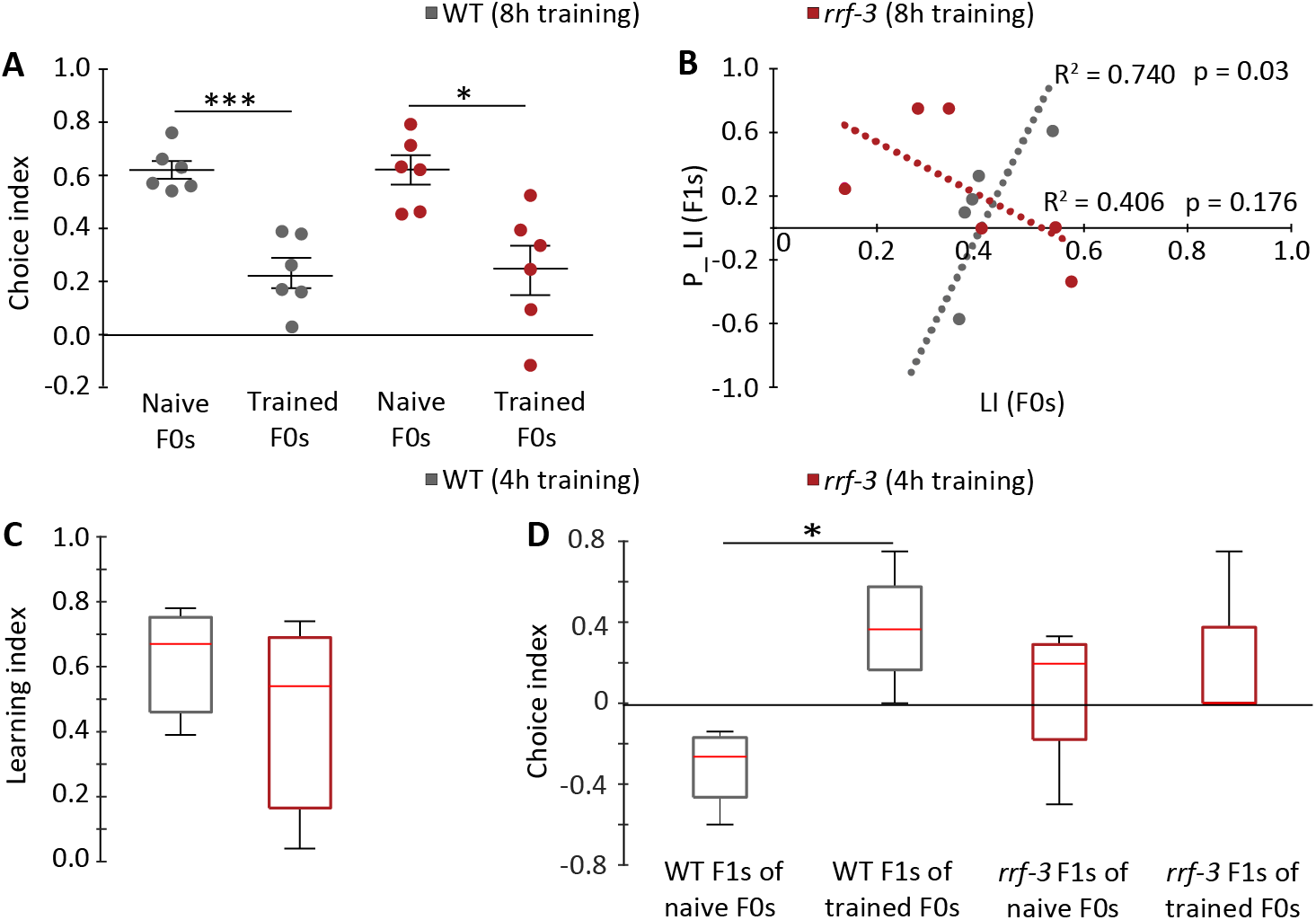
The RNA dependent RNA polymerase RRF-3 regulates the parental experience-dependent olfactory learning of PA14. **(A, B)** Training with PA14 for 8 hours significantly decreased the preference for the smell of PA14, which is indicated by a positive choice index, in the F0 wild-type (WT) and the F0 *rrf-3(pk1426)* mutant animals (**A**, n = 6 learning assays for wild type (143 animals) and the *rrf-3* mutant (144 animals); two-tailed Student’s t-test, *** p < 0.001 and * p < 0.05, Mean ± SEM); however, while wild-type animals displayed a strong correlation between the parental-experience dependent learning index (P_LI) of F1s with the learning index (LI) of F0s, the mutation in the *rrf-3(pk1426)* animals abolished the correlation [**B**, n = 6 learning assays for F0 (143 WT animals and the 144 *rrf-3* animals) and n = 6 two-choice assays where 88 WT F1s and 92 *rrf-3* F1s were individually tested; P values denote the significance of the linear correlation coefficients]. **(C, D)** Training with PA14 for 4 hours induced similar aversive learning of PA14 in the F0 wild-type (WT) and the F0 *rrf-3(pk1426)* [**C**, n = 3 learning assays for WT (36 animals) and *rrf-3* (36 animals), p = 0.71]; however, while the wild-type F1 of the trained F0s displayed a significantly higher preference for PA14 than the wild-type F1s of the naive F0s in the two-choice assay, the *rrf-3(pk1426)* F1s of the trained F0s and the *rrf-3(pk1426)* F1s of the naive F0s displayed similar preference (**D**, n = 4 two-choice assays where 48 WT F1s and 51 *rrf-3* F1s were individually tested, * p = 0.0286 for WT and p = 0.91 for *rrf-3)*. Wilcoxon rank-sum test because the data in **C** and **D** are not normally distributed, the box plots show the median, the first and the third quantile, 1.5 IQR.

We were intrigued by the bidirectional effect of the parental experience with PA14 on the F1’s response to PA14. Previous studies show that although adult worms reduce their preference for the smell of PA14 after training, they still prefer the PA14 smell more than non-food smells^5^, indicating that PA14 represents as a less preferred food source to the trained F0s. Thus, we tested the possibility that training F0s with a shorter duration, which reduces the infection of PA14 to the F0s, induces a preference for PA14 in the F1s. Interestingly, we found that shortening the training of adult F0s to 4 hours generated aversive learning of PA14 in F0s (Figure 3C) and induced a significant increase in the preference for PA14 in the F1s of the trained F0s in comparison with the F1s of the naive F0s (Figure 3D). Furthermore, we found that although the *rrf-3(pk1426)* F0s were normal in learning of PA14 after 4-hour training (Figure 3C), their F1s were defective in the induced preference for PA14, indicated by the similar preference displayed by the *rrf-3(pk1426)* F1s of the trained and the naive mothers (Figure 3D). These results further demonstrate that the parental experience can generate the appetitive and the aversive effects on the olfactory response of PA14 in the progeny and that both of the intergenerational effects are regulated by the endogenous RNAi pathway.

Previously, we show that the adult *C. elegans* learns to avoid the smell of pathogenic bacteria, such as the *Pseudomonas aeruginosa* strain PA14, after feeding on the pathogen for 4-8 hours^4,5^. Meanwhile, it is shown that exposing the L1-stage larvae worms to PA14 generates the avoidance of PA14 when the animals reach the adult stage and the imprinting of PA14 requires 12-hour training of the L1 larvae^22^. In this study we identify a new form of adaptive behavior where briefly training the adult worms with PA14 alters the olfactory response to PA14 in their progeny. Together, these results reveal versatile behavioral responses that *C. elegans* uses to interact with pathogenic bacteria that the worm frequently encounters in its habitat.

Unexpectedly, the experience with PA14 in the adults can regulate the olfactory response of their progeny bidirectionally. While the strong learning of the adult mothers leads to the avoidance of PA14 in the progeny, weak learning of the mothers induces attractive response to PA14 in their progeny. We found that training the mothers for 4 hours, the minimal amount of training needed to induce significant aversive learning of PA14 in the adults^5^, significantly increases the progeny’s preference for the smell of PA14, suggesting the mother-to-offspring transmission of beneficial information of PA14. The smell of PA14 is attractive to the worms that never ingest PA14; even after training, the PA14-trained adult worms that display a reduced preference for the PA14 smell still prefer the smell of PA14 more than the non-food odorants^5^. The bacteria of the *Pseudomonas* genus are highly abundant in the habits of *C. elegans* with many isolates being beneficial for the development and the reproduction of the worm^13^. Thus, it is conceivable that the parental experience transmits information of food and pathogenesis of PA14 to the progeny and the resulting behavioral effect is regulated by the extent of the experience.

In many animals, the parental experience with the environmental conditions, especially those associated with food and danger, regulates the development, physiology and behavior of the offspring and the effect can last for various amounts of time^1,3,23–29^. In *C. elegans*, the insulin-like signaling acts in the soma or the germline of the mothers to regulate the development and the stress responses in response to the parental experience with food deprivation or osmotic stresses, respectively^26, 27^. However, in most cases, especially when the parental experience generates adaptive changes in the same behavioral modality in both the mothers and the progeny, the signaling mechanism underlying the transmission of the parental experience is largely unknown. In *C. elegans*, it is shown that the pathways of the endogenous small interfering RNAs (endo-siRNAs) can regulate the gene expression and the function of the nervous system in response to the experience within the same generation^30,31^. In addition, it is shown that the production of endo-siRNAs is regulated by prolonged starvation and the altered siRNA profile persists for several generations^20^. In this study, we show that the RRF-3/RdRP^21^, which is critically required for the biogenesis of endo-siRNAs that are produced in the soma or maternally inherited^6,7^, specifically regulates the parental-experience dependent changes in the olfactory response to the pathogenic bacteria PA14, revealing a new function of the endo-siRNAs in regulating intergenerational adaptive behavioral responses.

## Methods

### Strains

*C. elegans* hermaphrodites were used in this study and cultivated using the standard conditions^16^. The strains used in this study include: N2 (Bristol), ZC2834 *rrf-3(pk1426)* II and CX4998 kyIs 140 I; *nsy-1(k397)* II.

### The aversive olfactory training with PA14 in F0s and the cultivation of the F1s

The aversive olfactory training was preformed mainly as previously described^4^ with minor modifications. To prepare the training plates, individual OP50 colonies or PA14 colonies were used to inoculate 50 mL nematode growth medium (NGM, 2.5g/L Bacto Peptone, 3.0g/L NaCl, 1mM CaCl2, 1mM MgSO4, 25mM KPO4 pH6.0), which were cultivated at 27°c for overnight. 800 mL (for 4h training protocol) or 400 mL (for 8h training protocol) OP50 or PA14 culture was spread onto each NGM plate, which was incubated at 27°c for 2 days to prepare the naive control and the training plates, respectively. To preform training, the first-day adult hermaphrodites cultivated under the standard conditions with OP50 as food were transferred to the control or the training plate, respectively, and kept at 20°c for 4 or 8 hours as specifically described. By the end of the training, some of the naive control and the trained F0 worms were randomly picked to measure the olfactory choice and learning in the automated assay and the rest of the F0s worms were collected from the plates with S-basal medium and passed through a cell strainer 40 micrometer nylon filter (Falcon) to remove laid eggs. Worms were then treated with a bleach solution to isolate the F1 embryos, which were hatched and cultivated under the standard conditions until the adult stage.

### Olfactory preference assay using the automated olfactory assay

The automated assay that quantifies the olfactory preference in individual animals were conducted as previously described^5^. Briefly, by the end of the training, the naive and trained F0s were washed with buffer and individually placed in the droplets of 2 μL NMG buffer (1mM CaCl2, 1mM MgSO4, 25mM KPO4 pH6.0) in an enclosed chamber and subjected to alternating airstreams that were saturated with the smell of OP50 or PA14 by blowing clean air through freshly generated bacterial cultures. Each olfactory stimulation lasted for 30 second and each assay contains 12 cycles of stimulation. The locomotion of the worms is video recorded and the large body bends of the tested worms were identified with machine-vision softwares. Because large body bends are followed by reorientation, a higher rate of the large body bends indicates a lower preference to the tested airstream^5,32^. The choice index (CI) is defined as the body bends evoked by OP50 smell minus the body bends evoked by PA14 smell and normalized by the total body bends [CI = (bends evoked by OP50 - bends evoked by PA14)/(bends evoked by OP50 + bends evoked by PA14)] and the learning index (LI) is defined as the CI of the naive worms minus CI of the trained worms (LI = CI of naive worms – CI of trained worms). A positive LI indicates learned avoidance of PA14.

### Olfactory preference assay using the two-choice assay

The two-choice assay is similar to the one described in Zhang et al., 2005, except for several modifications. To measure the olfactory preference for bacteria, a drop of 5 μL supernatant of OP50 culture and a drop of 5 μL PA14 culture were put 2cm apart on a 6 cm NMG plate. In each assay, one worm was placed on a plate equidistant to the two drops of the bacteria culture supernatant and allowed to crawl to the preferred stimulus. The movement and the choice of each worm were recorded and later analyzed. The worms that did not make choice after 10 minutes are counted in the total number. The parental-experience dependent choice index (P_CI) in the two-choice assay is defined as the number of worms that choose PA14 minus the number of worms that choose OP50 normalized by the total number of worms tested [P_CI = (number of worms that choose PA14 - number of worms that choose OP50)/total number of worms] and the parental-experience dependent learning index (P_LI) for the two-choice assay is defined as the P_CI of F1s of naive F0s minus P_CI of F1s of trained F0s (P_LI = P_CI of F1s of naive F0s – P_CI of F1s of trained F0s). A positive P_LI indicates increased avoidance of PA14 in F1 induced by the parental experience with PA14.

### Slow killing assay

The slow killing assay was performed essentially as previously described^14^. 200 μL freshly prepared PA14 culture incubated at 27°c in the Luria-Bertani (LB) medium overnight were spread into a 4cm diameter circle on a 6cm NMG plate and incubated at 37°c for 24 hours and then left at room temperature for another 24 hours before the assay. 20 F1 young adult hermaphrodites cultivated under the standard conditions (Figure 1A) were transferred onto each slow killing plate and kept at 20°c. The dead and the alive worms were counted at the specific time points as shown in Figure 2 over a course of 50 hours.

### The analysis of parental-experience dependent learning index

To separate the experiments in which F0s exhibited high learning indexes from the experiments in which F0s exhibited low learning indexes, we tested for a significant linear correlation between the learning index (LI) of F0s and the parental-experience dependent learning index of F1s (P_LI). We found that these 2 variables are linearly correlated as shown in Figure 1B. We then used the point where the line of linear regression intersects with the x-axis (LI of F0) to separate the experiments into two groups. The experiments in which the LI of F0 is larger than the point of intersection are included in the group in which F0s exhibited a high level of PA14 learning; conversely, the experiments in which the LI of F0 is smaller than the point of intersection are included in the group in which F0s exhibited a low level of PA14 learning.

### Quantification of the body size and the chemotactic movements

The body size and the parameters of the chemotactic movements in the two-choice assays were quantified by analyzing the recorded worms with the Wormlab tracker (https://www.mbfbioscience.com/wormlab).

## Statistics

The statistical methods, sample size and the number of the replicates are indicated in the legend of each figure.

## Data Availability

All the data are available upon request. Figure 1-3 and Extended Figure 1 and 2 are generated based on raw data.

## Code availability

The method used for the automated assay has been previously published^5^ and the code is freely available upon request. The software used to analyze F1 chemotaxis is available at https://www.mbfbioscience.com/wormlab.

## Acknowledgements

We thank *Caenorhabditis Genetics Center*, which is funded by National Institutes of Health - Office of Research Infrastructure Programs (P40 OD010440), for strains. Y.Z. is funded by National Institutes of Health.

## Author contributions

A.P., X.G., K.K. and Y.Z. designed the experiments, interpreted the results and wrote the manuscript. A.P., X.G. and K.K. performed the experiments and analyzed the data.

## Competing Interests

The authors declare no competing interests.

---

**Extended Figure 1.**
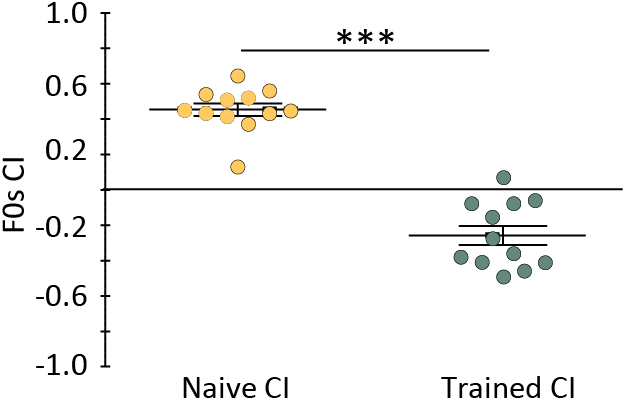
Training with PA14 for 8 hours strongly reduced the preference for the smell of PA14. The preference of the naive and trained F0 worms were measured in the automated assay. Choice index of F0 (CI) = (bends evoked by OP50 - bends evoked by PA14)/(bends evoked by OP50 + bends evoked by PA14); therefore, a positive choice index indicates a preference for the PA14 smell. n = 12 experiments (274 animals), two-tailed Student’s t-test, Mean ± SEM, *** p < 0.001.

**Extended Figure 2.**
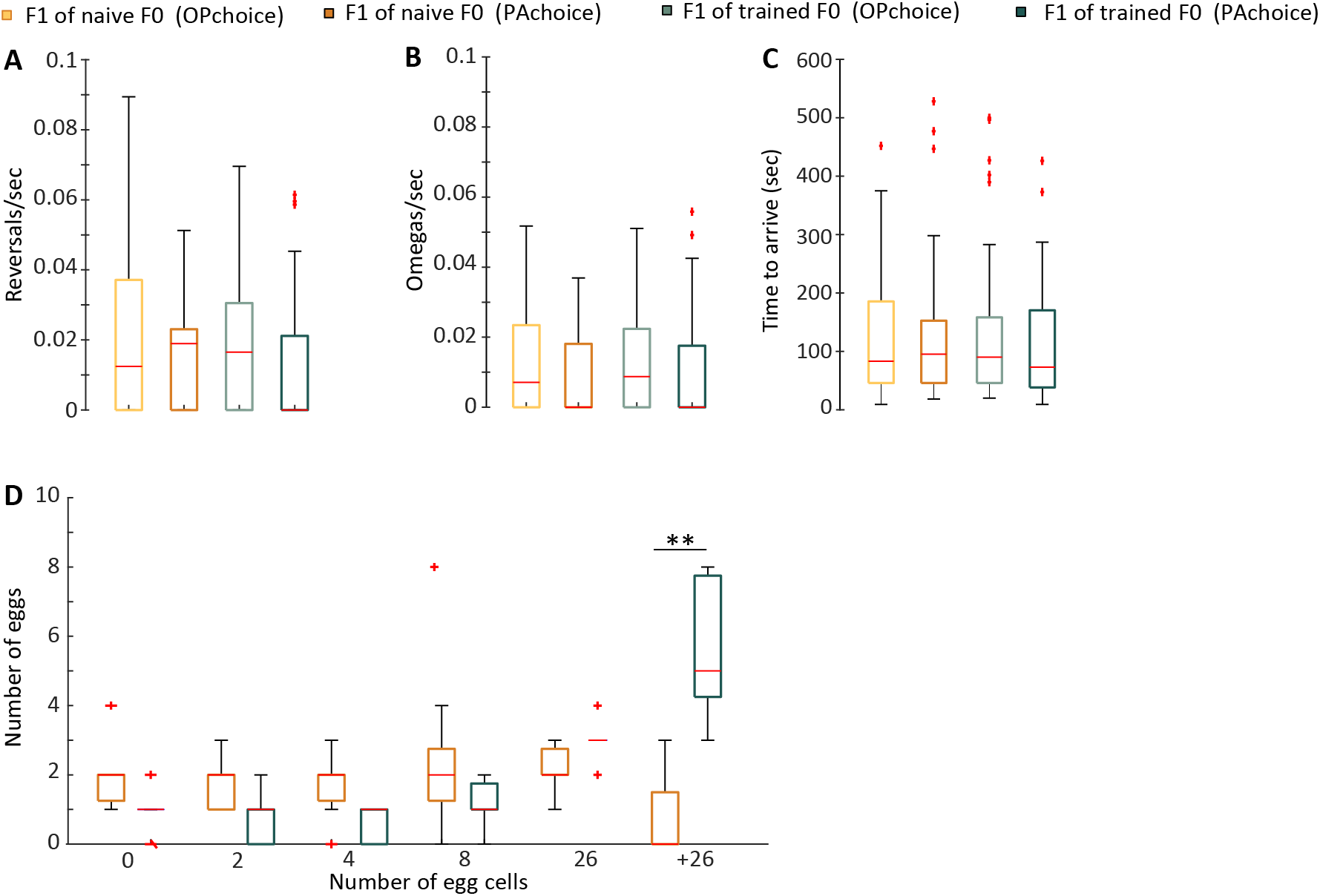
The parental experience with PA14 regulates the olfactory behavior and the development of the F1 progeny. **(A, B, C)** The F1s of the trained F0s and the F1s of the naive F0s do not differ in the rate of reversals **(A)** or omega bends (**B**, the large body bends resembling the Greek letter Ω that lead to reorientation) during chemotactic movements towards a drop of bacteria culture in the two-choice assay, or in the time needed to reach the target **(C)**. The box plots show the median, the first and the third quantile, 1.5 IQR and outliers; the number of worms for each group is the same as stated for Figure **2B**, Wilcoxon rank-sum test with correction for multiple comparison was used because the data is not normally distributed. **(D)** The eggs inside the bodies of the trained F0s are older than the eggs inside the naive F0s. The number of cells in each egg was counted. The box plots show the median, the first and the third quantile, 1.5 IQR and outliers; all the eggs in n = 7 worms per treatment were counted, Wilcoxon rank-sum test with correction for multiple comparison was used because the data is not normally distributed, ** p < 0.01.

